# Increased prefrontal top-down control in older adults predicts motor performance and age-group association

**DOI:** 10.1101/2020.12.09.417568

**Authors:** Philipp Alexander Loehrer, Felix Sebastian Nettersheim, Carina Oehrn, Fabienne Homberg, Marc Tittgemeyer, Lars Timmermann, Immo Weber

**Affiliations:** Department of Neurology, University Hospital Gieβen and Marburg, Marburg, Germany; Department of Neurology, Philipps-University Marburg, Marburg, Germany; Department of Cardiology, University Hospital Cologne, Cologne, Germany; Center for Mind, Brain and Behavior (CMBB), Philipps-University Marburg, Marburg, Germany; Max Planck Institute for Metabolism Research, Cologne, Germany; Cologne Cluster of Excellence in Cellular Stress and Aging-Associated Disease (CECAD), Cologne, Germany

**Author notes:** <u>Corresponding author</u>: Philipp Loehrer, Department of Neurology, University Hospital Marburg, Baldinger Str., 35043, Marburg.

## Abstract

Bimanual motor control declines during ageing, affecting the ability of older adults to maintain independence. An important underlying factor is cortical atrophy, particularly affecting frontal and parietal areas in older adults. As these regions and their interplay are highly involved in bimanual motor preparation, we investigated age-related connectivity changes between prefrontal and premotor areas of young and older adults during the preparatory phase of complex bimanual movements using high-density electroencephalography. Generative modelling showed that excitatory interhemispheric prefrontal to premotor coupling in older adults predicted age-group affiliation and was associated with poor motor-performance. In contrast, excitatory intrahemispheric prefrontal to premotor coupling enabled older adults to maintain motor-performance at the cost of lower movement speed. Our results disentangle the complex interplay in the prefrontal-premotor network during movement preparation underlying reduced bimanual control and the well-known speed-accuracy trade-off seen in older adults.

## 1. Introduction

Physiological ageing is associated with a decline in fine motor control^1^. Deficits in motor performance arising with age comprise coordination problems, reduced movement variability, as well as an overall slowing of movements^1^. Although the ability of older adults (OA) to perform learned and frequently practiced actions as well as automatized movements is not impaired^2^, older adults are often overchallenged by complex tasks that require a high amount of flexibility^3^. The decline in motor control in OA becomes particularly apparent during the performance of bimanual movements affecting activities of daily living and the ability to maintain independence^1, 4^. Hence, in an ageing society, it is of great interest to understand the mechanisms of physiological ageing and associated deterioration of motor functions.

An important underlying factor of age-related motor deficits is a loss of brain volume^1^. Several studies have linked reduced gray matter volume as well as decreased white matter integrity in OA to reduced motor performance (for a review see Seidler et al.^1^). During ageing, frontal areas, including the prefrontal cortex, as well as parietal areas seem to be particularly susceptible to atrophy^1^. Based upon these findings, an initial hypothesis within the cognitive literature proposed that a reduced commitment of atrophied brain regions underlies the performance deficits in OA^1^. To study age-related changes in functional brain recruitment patterns non-invasively, the methods of functional magnetic resonance imaging (fMRI) and electroencephalography (EEG) have been utilized. During processes of cognition and memory retrieval, OA indeed show a pattern of reduced cortical activation^1^. In contrast, when engaging in executive functions and working memory tasks, OA show greater activation in prefrontal and parietal areas^1^. These findings have been extended into the literature on motor control. Here, over-activation was particularly present in frontal^5, 6, 7^ and parietal areas^6, 7, 8^ during unimanual^8, 9^ and bimanual motor tasks^6, 7^. Over time, two theories have emerged to explain over-activation in the ageing brain. The de-differentiation hypothesis states that the increased activation pattern represents a non-selective and inefficient recruitment of brain areas^10^. In contrast, the more frequently invoked compensation hypothesis states that over-activation compensates for a reduction in the brain’s structural integrity and is associated with better performance^9^. During motor tasks, the pattern of over-activation was predominantly associated with increased motor performance suggesting a compensatory mechanism. Therefore, over-activation of higher-level prefrontal and sensorimotor cortical areas in older adults is proposed to represent an increased reliance on cognitive control and sensory information processing to match the motor performance of young adults (YA)^1^. This movement-related pattern of over-activation in OA is accompanied by distinct connectivity changes in the motor system^8, 11, 12, 13^. Overall, an increase in connectivity metrics, such as connectivity strength and network efficiency, analyzed through generative modelling, was associated with ageing and – predominantly – preserved motor function^11, 12^. Particularly, OA showed increased prefrontal influences on the motor system while parietal influences decreased during movement execution^12^. Furthermore, distinct connections exhibited reduced connection strengths in OA during the execution of unimanual finger tapping (left PFC to left premotor cortex)^8^ and unilateral foot moving (right M1 to right premotor cortex)^13^. These findings support the notion of increased reliance on cognitive control during movement execution in older adults but also demonstrate that the influence of age depends on the connection studied.

Since areas primarily affected by age-related atrophy and subsequent changes in activity and connectivity are highly involved in unimanual and particularly bimanual movement preparation, age-related changes during the preparatory phase of movements are of particular interest. In this regard, Berchicci and colleagues could demonstrate that OA prepared simple and complex unimanual finger movements with higher prefrontal activation to reach accuracy levels comparable to those of YA^14^, suggesting that OA rely on increased cognitive control during movement preparation also. Furthermore, the lateralized activation seen over the hemisphere contralateral to the moving hand during the preparation of a movement in YA is diminished in OA. Rather, OA show an increased, i.e. more positive, amplitude of distinct event-related potentials (lateralized readiness potential, P300, contingent negative variation)^15, 16^. Here, Hinder and colleagues showed that interhemispheric inhibition (IHI) between premotor and motor areas is reduced (i.e. more facilitatory) during preparation of unimanual finger movements in OA^17^, supporting the idea that increased amplitudes in OA arise from a reduction in IHI^18^. On the other hand, strictly timed interhemispheric inhibition during movement preparation is essential for the correct execution of bimanual movements^19^. Therefore, age-related changes of interhemispheric connectivity during movement preparation could be an important underlying factor of reduced bimanual control in older adults.

To date, the influence of age on the interplay between bilateral prefrontal and premotor areas during the preparation of bimanual movements remains to be elucidated. Furthermore, evidence regarding the individual electrophysiological differences between older adults with preserved and reduced bimanual coordination is lacking. Based on the findings named above we hypothesized that during bimanual movement preparation (1) OA exhibit increased connection strengths compared to young adults, (2) increased connectivity strengths are related to differences in behavioural parameters, and (3) older adults with preserved bimanual coordination show connectivity patterns resembling those of young adults. To test these hypotheses, we assessed effective connectivity in young and older adults during the preparatory phase of bimanual movements. Participants performed temporally coupled and spatially uncoupled movements while a 128-channel electroencephalography (EEG) was recorded. To infer connectivity between areas highly involved in bimanual movement preparation including prefrontal cortex (PFC), lateral premotor cortex (lPM), and supplementary motor area (SMA), we used Dynamic Causal Modelling for evoked responses (DCM for ERP)^20^. DCM for ERP is a statistical approach employing neuronal mass models to infer effective connectivity between distinct neuronal sources based on measured EEG responses^21, 22^. To delineate differences between age groups and relate effective connectivity during movement preparation to behavioural parameters, we employed the Parametric Empirical Bayes (PEB) framework^23, 24^.

## 2. Results

### 2.1 Behavioral results

We explored the overall performance difference between older adults (n=20) and young adults (n=19) employing two-tailed independent samples t-tests. Here, older adults made more mistakes compared to young adults, however, this difference was not significant (t(37) = 1.278, p = .209, Cohen’s d = .41, older adults: 28.33±1.35 (mean±SEM), 95% CI [25.67-30.98], young adults: 22.87±1.19, 95% CI [20.54-25.19]). Furthermore, older adults needed more time to perform the task compared to young adults (t(37) = 3.452, p = .001, Cohen’s d = 1.11, older adults: 3.09±0.06, 95% CI [2.98-3.20], young adults: 2.34±0.04, 95% CI [2.26-2.42]). To evaluate differences in top performing older adults (n=10) compared to low performers (n=10) and young adults (n=19), we used a mixed design analysis of variance (ANOVA). For the variable *error rate*, an overall between-subject effect of “group” (F(2,36) = 7.658, p = .002, Cohen’s f: .65) was revealed. Post hoc Tukey tests demonstrated that old – low performers made more mistakes compared to old – top performers (p = .002, Cohen’s d = 1.18, old – low performer: 38.86±1.96, 95% CI [35.01-42.70], old – top performer: 17.8±1.29, 95% CI [15.28-20.32]) and compared to young adults (p = .009, Cohen’s d = .83). No difference could be observed between old – top performers and young adults (p = .542, Cohen’s d = .28, Figure 1.A). Moreover, there was a trend toward a significant main effect of complexity (F(4, 142) = 2.38, p = .055, Cohen’s f: .26). No further main effects or interactions were revealed for *error rate* (all p > .108). Mixed design ANOVA of *performance time* similarly revealed an overall between-subject effect of “group” (F(2,36) = 7.837, p = .001, Cohen’s f = .66). Post hoc Tukey tests showed that old – low performers needed more time compared to young adults (p = .001, Cohen’s d = 1.48; old – low performer: 3.34±0.07, 95% CI [3.22-3.48]). No difference could be observed between old – top performers and young adults (p = .155, Cohen’s d = .65; old top – performer: 2.83±0.08, 95% CI [2.67-2.99]) as well as old – low performers and old – top performers (p = .195, Cohen’s d = .67, see Figure 1.B). There were main effects of hand (F(1, 36) = 14.765, p < .001, Cohen’s f: .64; subjects needed more time when the learned sequence was tapped with the right hand) and complexity (F(3.5, 126) = 11.212, p < .001, Cohen’s f: .56). Moreover, there was an interaction between the factors hand and complexity for *performance time* (F(4,146) = 2.892, p = .024, Cohen’s f: .28). No further significant main effects or interactions were revealed for *performance time* (all p > .085).

**Figure 1.**
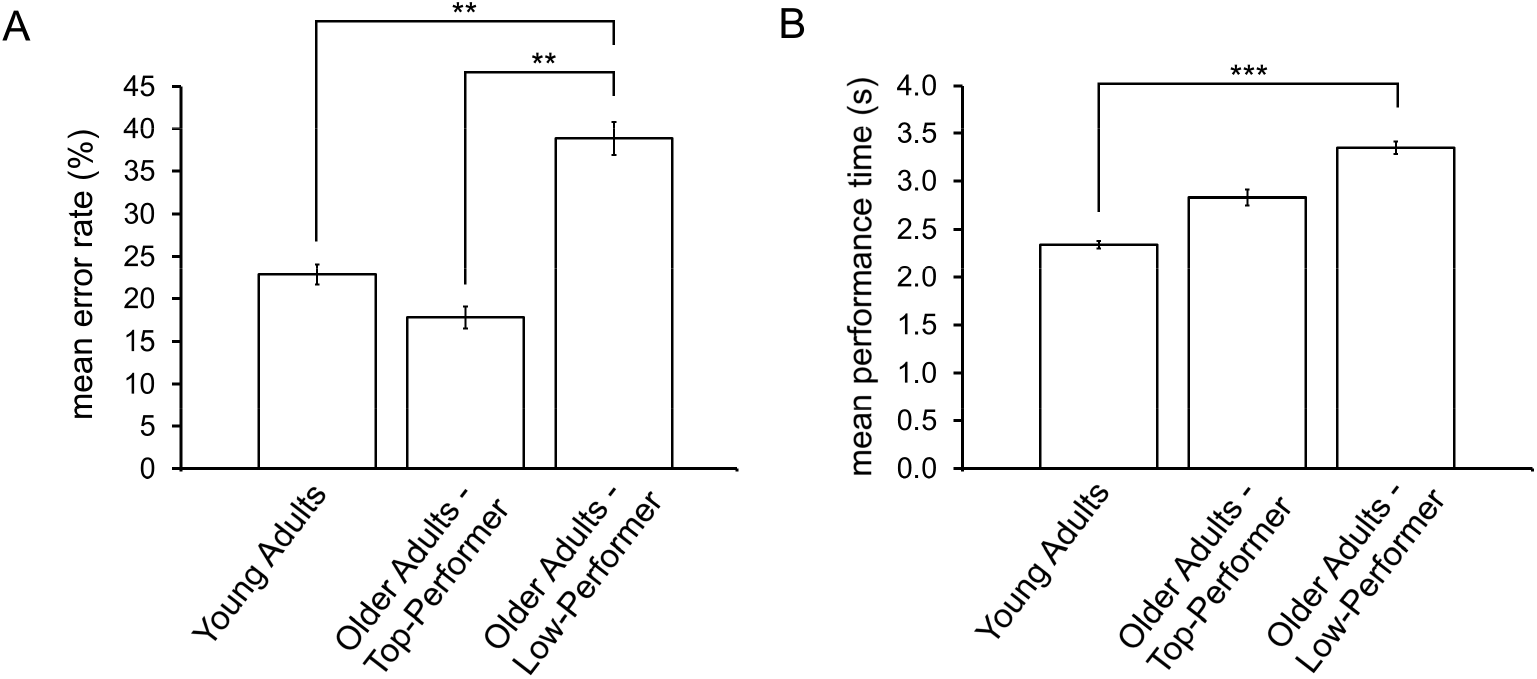
Behavioral differences between young and older adults. **A.** Mean error rates and standard error of the mean (SEM) for young (n=19) and older adults (n=20). Old – low performers (n=10) made more mistakes compared to young adults (p = .009; all post hoc tests are Tukey tests) and older adults in the top-performing group (n=10; p = .002). No difference was observed between YA and Old – top-performer (p = .542). **B.** Mean performance time and SEM for young and older adults. Old – low-performers needed more time to complete the task compared to YA (p < .001). No difference could be observed between old – top performers and YA (p = .155) as well as old – top performers and old – low performers (p = .195). Asterisks indicate significant differences (** p ≤ .01, *** p ≤ .001).

### 2.2 Event Related Potentials - Sensor Space Analysis

To delineate differences in neural responses during movement preparation between older and young adults we investigated event related potentials time locked to the “go-stimulus” employing a one-sided permutation test. In both conditions older adults (n=20) exhibited more positive ERPs than young adults (n=19). If the learned sequence was tapped with the right hand, statistical analysis of ERPs revealed a group-difference between 517 and 758 ms post-stimulus (p = .003, Cohen’s d = 1.42). Furthermore, we found differences in ERPs between 793 and 1000 ms post-stimulus intermittently. Differences were located in the sensors above the right hemisphere over the postcentral sulcus and the angular gyrus (Figure 2.A and C). If the learned rule was executed with the left hand, however, differences were found in the left hemisphere, more anteriorly above the frontal gyrus in a latency range of 313 to 570 ms (p = .043, Cohen’s d = 1.22, Figure 2.B and D). ERP-analysis of YA compared to OA stratified by performance yielded similar results (cf. Supplementary Figure 1 and 2). Here, differences between YA and OA – top performers (n=10) showed differences in the same latency ranges over the same areas, whereas differences between YA and OA – low performers (n=10) were non-significant for condition 2 and showed a trend toward significance for condition 1.

**Figure 2:**
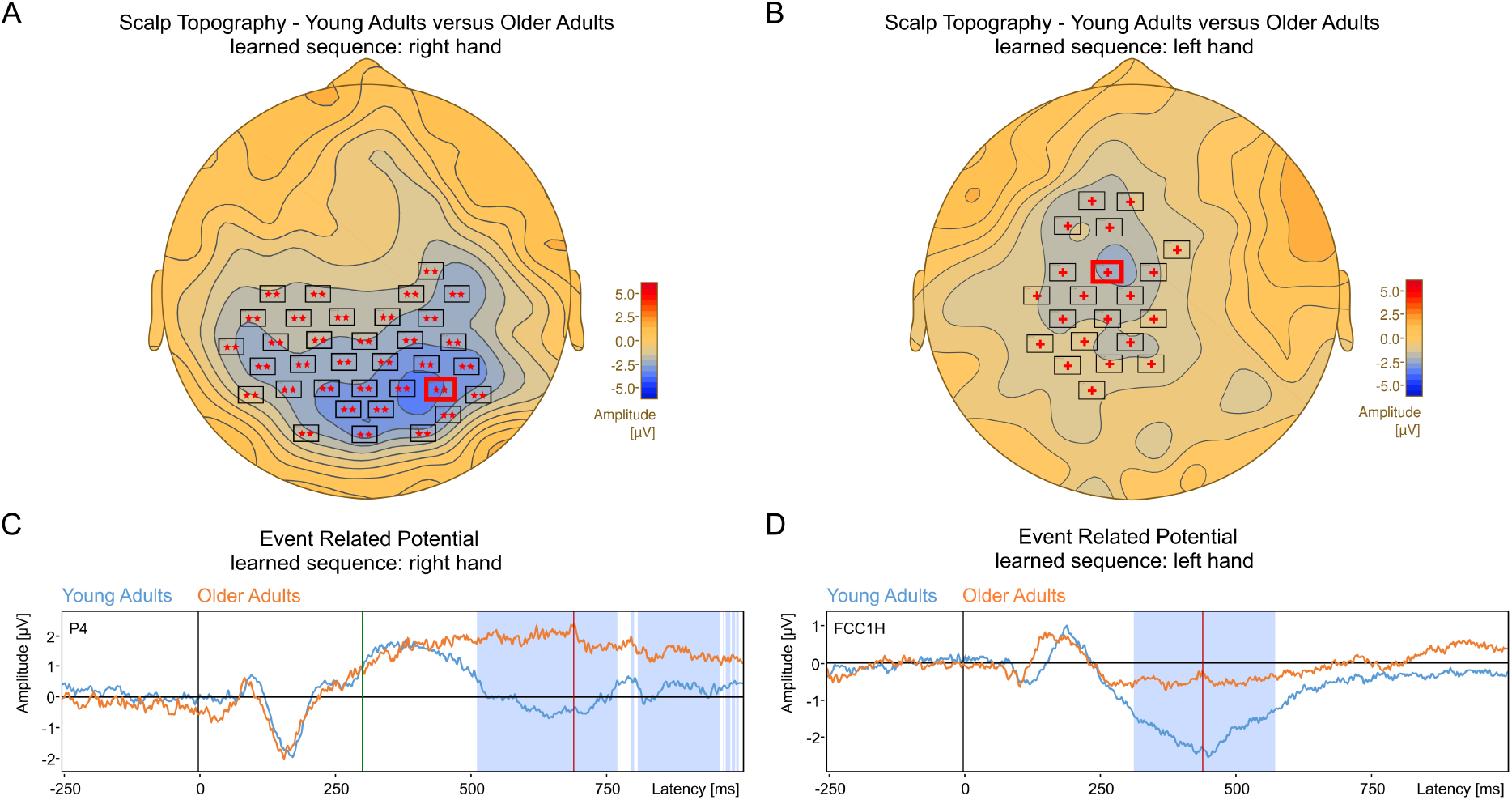
Scalp topography of EEG sensors showing significant differences between grand-mean ERPs for young and older adults. **A.** Scalp topography plotted for correctly performed trials at 688 ms after stimulus onset when the learned rule was performed with the right hand and the new number sequence with the left hand. The cluster with a p-value = .003 is denoted with two starlets (one-sided permutation test, YA: n=19, OA: n=20). The red line shows the time point for which the scalp topography is plotted. **B.** Scalp topography for correctly performed trials when the learned rule was performed with the left hand and the new number sequence with the right hand for 440 ms after stimulus onset. The cluster with a p-value of .043 is denoted with a cross (one-sided permutation test, YA: n=19, OA: n=20). The red line shows the time point for which the scalp topography is plotted. The dark green line denotes 300 ms post “go”-stimulus. **C.** ERP waveforms averaged time-locked to the “go”-stimulus: Grand-mean ERP of young (blue curve) and older (orange curve) adults recorded with electrode P4 (right hand rule). Significant differences are shaded in light blue color. **D.** ERP waveforms averaged time locked to the “go”-stimulus: Grand-mean ERP of young and older adults recorded with electrode FCC1H (left hand rule). Significant differences are shaded in light blue color.

In summary, differences were located on the hemisphere contralateral to the hand tapping the new number sequence in both conditions.

### 2.3 Dynamic Causal Modelling – Source Space Analysis

As no evidence from anatomical studies regarding lPM and SMA hierarchy exists, we performed a random effects Bayesian model comparison (BMC) between three fully connected models with either forward, backward, or lateral connections between lPM and SMA. BMC favored a model featuring forward connections from SMA to lPM and backward connections from lPM to SMA (Model Exceedance Probability: 0.70; see Supplementary Figure 3). Ensuing PEB models were based on this model structure. Here, we took intrinsic as well as extrinsic connectivity strengths to the group level and used Bayesian Model Reduction (BMR) to exclude connections that did not contribute to the model evidence. Commonalities between participants as well as differences due to age-group, performance-group, *error rate*, and *performance time* were determined for parameters of A- and B-Matrix separately. Here, mean connectivity parameter estimates were derived by averaging the parameters across PEB models weighed by the models’ posterior probability, a process called Bayesian Model Averaging.

#### 2.3.1 A-Matrix Parameters – Commonalities

Bayesian model averages of the A-Matrix parameters (average effective connectivity across conditions) that were in common over subjects showed a consistent inhibitory connectivity profile between premotor and prefrontal areas (Figure 3.A-B). Here, forward connections from left lPM to left PFC and right lPM to right PFC were of inhibitory nature. Similarly, forward connections from SMA to left and right PFC were inhibitory. Spearman correlation analysis revealed that weaker inhibitory influence of lateral premotor areas on prefrontal Cortex positively correlated with higher *error rates* (left lPM to left PFC: YA: *r* = .99, *p* < .001, OA: *r* = .98, *p* < .001; right lPM to right PFC: YA: *r* = .90, *p* = < .001, OA: *r* = -.21, *p* = .45, Figure 3.C-D). Weaker inhibitory influence of SMA on PFC, on the other hand, was correlated with higher performance time in young adults (SMA to left PFC: *r* = .57, *p* = .024; SMA to right PFC: *r* = .69, *p* = .004, Figure 3.E-F). In older adults, this association was not significant (SMA to left PFC: *r* = .03, *p* = .95; SMA to right PFC: *r* = .10, *p* = .78). When differentiating between old top- and low-performers, however, this association reached statistical significance at the .05-level (SMA to left PFC: old top-performer: *r* = .92, *p* = .001, old low-performer: *r* = .50, *p* = .22; SMA to right PFC: old top-performer: *r* = .98,*p* < .001, old low-performer: *r* = .84, *p* = .009, Figure 3.E-F).

**Figure 3.**
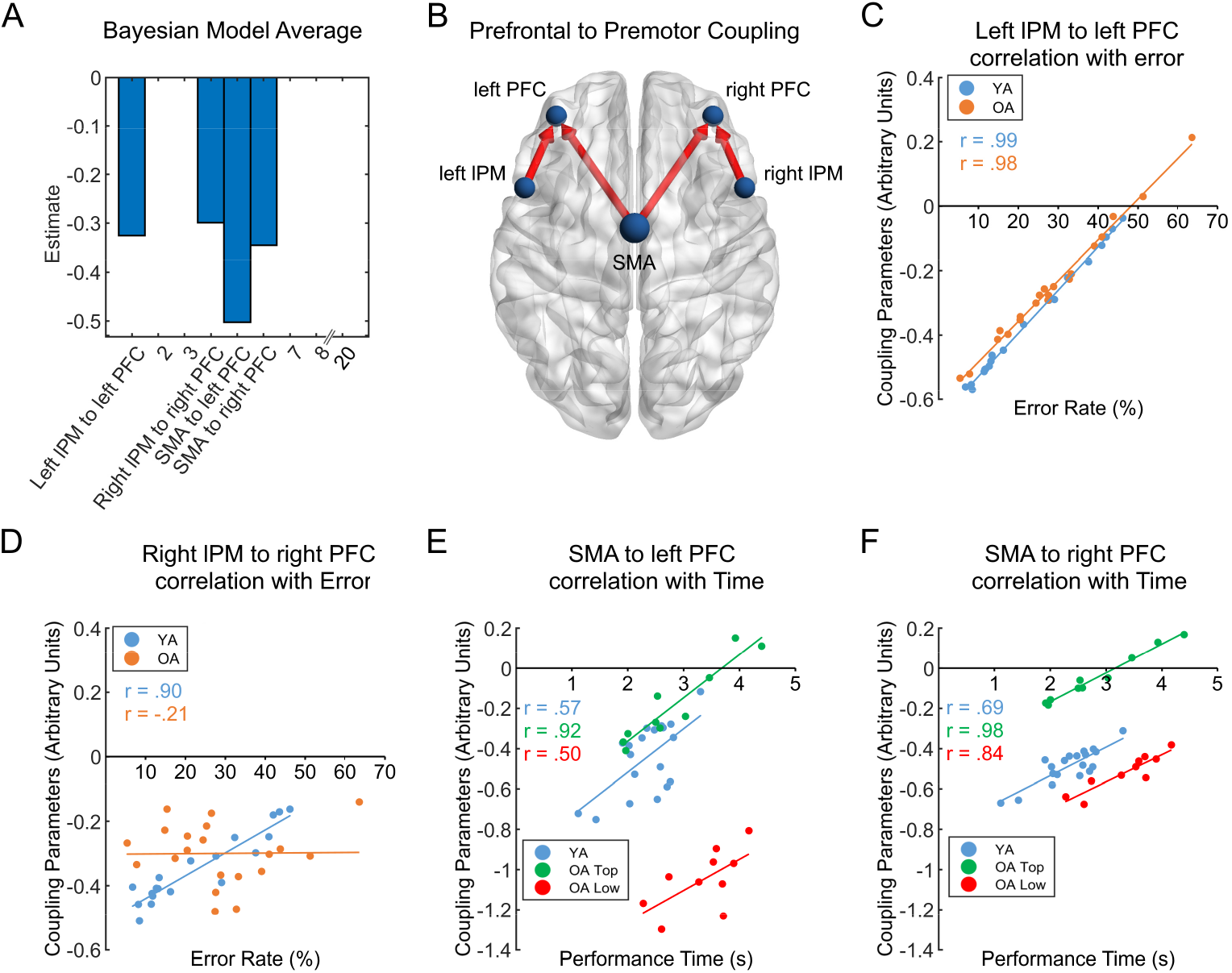
**A.** Bayesian model averages of the commonalities between groups thresholded at posterior probability >95% for clarity. The schematic in **B** illustrates the parameters that survived thresholding. Red arrows denote inhibitory connections. **C-D.** Error rates plotted against individual coupling parameters of lateral premotor (lPM) to prefrontal cortex (PFC) coupling (YA: young adults, n=19; OA: older adults, n=20). **E-F.** Performance time plotted against individual coupling parameters of supplementary motor area (SMA) to PFC coupling (OA Top: older adults – top performer (n=10), OA Low: older adults – low performer (n=10)). The schematic was visualized with BrainNet Viewer^25^.

#### 2.3.2 A-Matrix Parameters – Age-Group Differences

Bayesian model averages of parameters related to differences in age-group showed a consistent pattern of being more excitatory in older adults. Here, interhemispheric backward connections from left PFC to right lPM as well as from right PFC to left lPM were more excitatory in OA (Figure 4.A-B). Spearman correlation analysis revealed that a weaker inhibitory influence of left PFC on right lPM in YA and a stronger excitatory influence in OA correlated with higher *error rates* (left PFC to right lPM: YA: *r* = .85, *p* < .001, OA: *r* = .93, *p* < .001, Figure 4.D).

**Figure 4.**
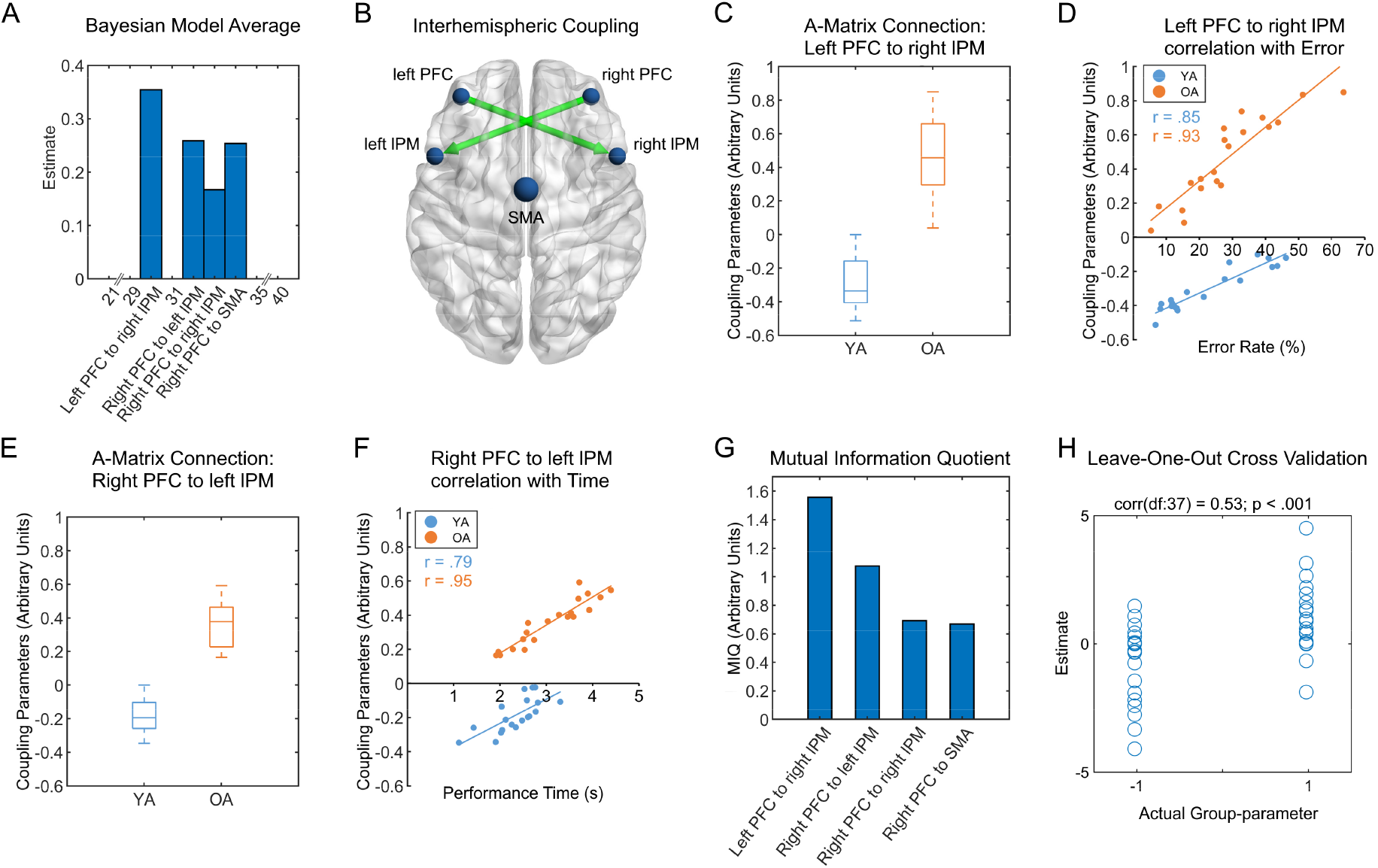
**A.** Bayesian model averages of the group differences thresholded at posterior probability >95% for clarity. The four parameters displayed are the only parameters that survived thresholding. The schematic in **B** illustrates the interhemispheric parameters that survived thresholding. Green arrows denote more excitatory connections in older adults compared to young adults. **C and E.** Box-plots for coupling parameters of connections left PFC to right lPM (**C**) and right PFC to left lPM (**E**, YA: young adults (n=19), OA: older adults (n=20)). Center line indicates the median, box limits represent upper and lower quartiles and whiskers indicate most extreme data points not considered outliers **D.** Error rates plotted against individual coupling parameters of left PFC to right lPM coupling. **F.** Performance time plotted against individual coupling parameters of right PFC to left lPM. **G.** Mutual information quotients for coupling parameters that survived thresholding. Interhemispheric connections were selected to reduce dimensionality and increase classification accuracy. **H.** Results of leave-one-out cross validation. Actual subject group-parameter (YA = −1, OA = 1) plotted against the expected value of the estimated group-parameter.

On the other hand, weaker inhibitory influence of right PFC on left lPM in young adults and a stronger excitatory influence in older adults correlated with higher *performance time* (right PFC to left lPM: YA: *r* = .79, *p* < .001, OA: *r* = .95, *p* < .001, Figure 4.F). A post-hoc ANOVA revealed that OA in the top-performing group had lower connection strengths compared to subjects in the low-performing group for both interhemispheric connections. Here, the connection strength from left PFC to right lPM was significantly lower (*p* < .001) and connection strength from right PFC to left lPM showed a trend toward significance for being lower in the top-performing group (*p* = .06). Thus, old top-performers had connectivity profiles more similar to young adults.

Furthermore, backward connections from right PFC to right lPM and from right PFC to SMA were more excitatory in older adults. Spearman correlation analysis revealed that a stronger inhibitory influence of right PFC on right lPM in young adults was associated with higher *error rates* (right PFC to right lPM: young adults: *r* = -.91, *p* <.001, Figure 5.D). In older adults, this association was not significant (old adults: *r* = -.35, *p* = .21). When differentiating between old top- and low-performers, however, the association between weaker excitatory influence and higher error rates reached statistical significance at the .05-level (old top-performer: *r* = -.84, *p* = .009, old low-performer: *r* = -.87, *p* = .006, Figure 5.D). Additionally, a weaker inhibitory influence of right PFC on SMA in young adults and a stronger excitatory influence in older adults was related to higher *performance time* (young adults: *r* = .87, *p* <.001, older adults: *r* = .89, *p* <.001, Figure 5.F). Here, no differences in connection strengths between top- and low-performing older adults could be revealed by the post-hoc ANOVA.

**Figure 5.**
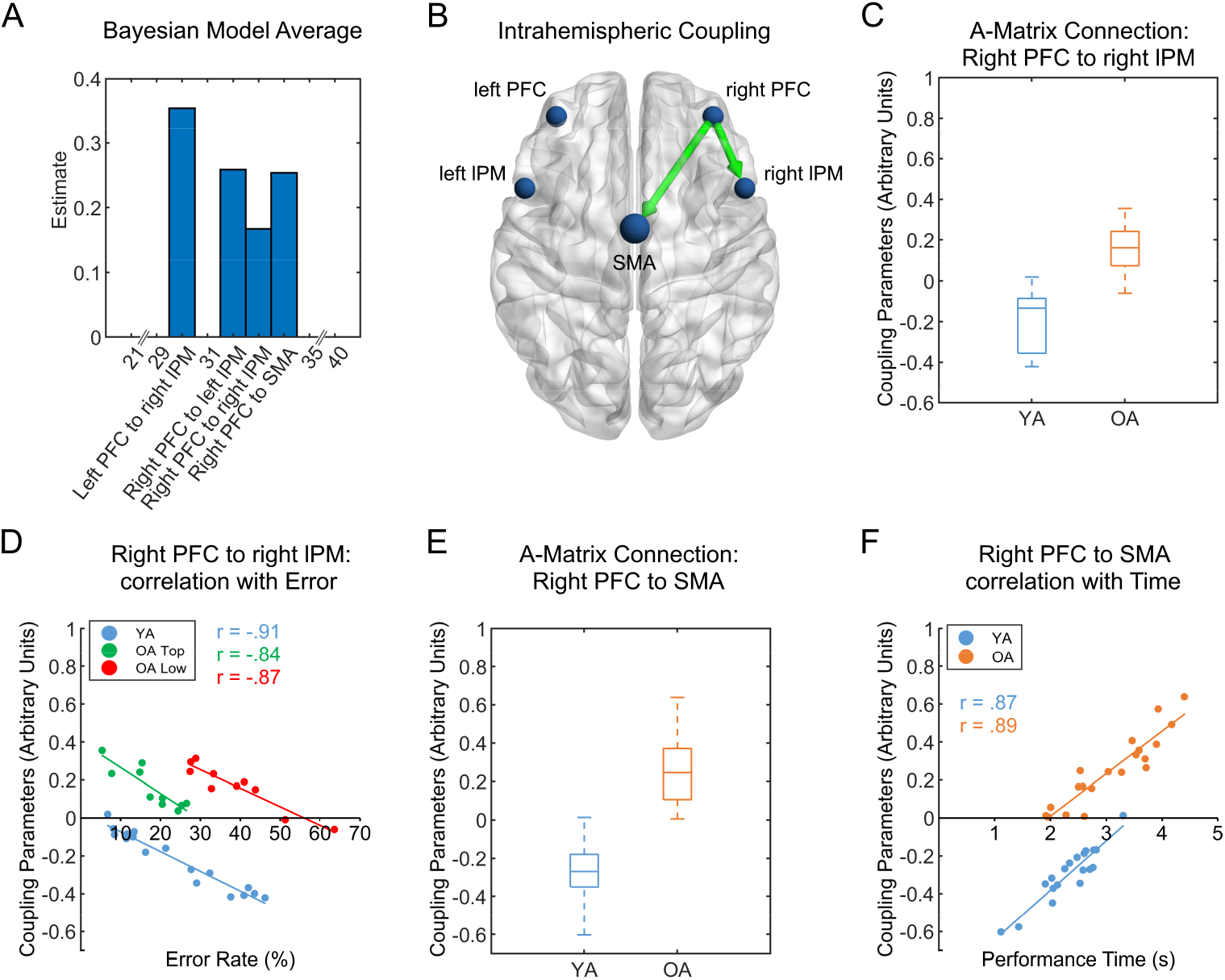
**A.** Bayesian model averages of the group differences thresholded at posterior probability >95% for clarity. The four parameters displayed are the only parameters that survived thresholding. The schematic in **B** illustrates the intrahemispheric parameters that survived thresholding. Green arrows denote more excitatory connections in older adults compared to young adults. **C and E.** Box-plots for coupling parameters of connections right PFC to right lPM (**C**) and right PFC to SMA (**E**, YA: young adults (n=19), OA: older adults (n=20)). Center line indicates the median, box limits represent upper and lower quartiles and whiskers indicate most extreme data points not considered outliers. **D.** Error rates plotted against individual coupling parameters of right PFC to right lPM coupling (OA Top: older adults – top performer (n=10), OA Low: older adults – low performer (n=10)). **F.** Performance time plotted against individual coupling parameters of right PFC to SMA coupling (YA: young adults (n=19), OA: older adults (n=20)).

#### 2.3.3 B-Matrix Parameters – Commonalities and Age Group Differences

To evaluate the influence of age on the differences between hands, we used the PEB framework to model the B-Matrix parameters. Here, Bayesian model averages of parameters related to commonalities between subjects showed that the backward connection from left PFC to SMA was decreased in condition 1 (learned sequence tapped with right hand) compared to condition 2 (learned sequence tapped with left hand). Furthermore, connections from right lPM to SMA and from left lPM to left PFC were increased, whereas the influence of left lPM on itself was reduced in condition 1 compared to condition 2. Spearman correlation analysis revealed that a larger difference in PFC to SMA connectivity strength between both conditions correlated with a larger difference in *performance time* in all subjects (left PFC to SMA: *r* = .70, *p* < .001). Weaker PFC to SMA connectivity in condition 1 was hence associated with an increased *performance time*. There were no other correlations between connectivity parameters related to commonalities and performance differences. Furthermore, there were no difference in the B-Matrix parameters due to age-group.

### 2.4 Mutual Information Quotient and Leave-One-Out Cross Validation

To assess the predictive validity of the A-matrix connectivity parameters, we employed leave-one-out cross validation. MIQ feature selection algorithm selected both interhemispheric connections (Figure 4.G). Thus, we used these connections for leave-one-out-analysis (LOO). Here, the estimated effect sizes of interhemispheric connections were large enough to predict age-group affiliation (Pearson’s correlation coefficient: 0.53, *p* < .001, Figure 4.H).

## 3. Discussion

Using Dynamic Causal Modelling for evoked responses and Bayesian Model Averaging, we show for the first time that excitatory interhemispheric top-down control from prefrontal to lateral premotor cortex predicts age-group association and motor performance. Furthermore, we report that right-hemispheric top-down control from prefrontal cortex to premotor areas underlies the commonly observed finding of preserved accuracy at the cost of increased performance time in older adults. Our results disentangle the complex interplay between prefrontal and premotor areas during movement preparation and describe distinct connectivity profiles that predict poor as well as preserved motor performance in older adults.

### 3.1 Preserved accuracy at the cost of increased performance time in older adults

It is a common finding that older adults can match the accuracy of young adults in simple tasks^11, 26^. When task complexity rises, however, OA have to compensate for the increased task demands at the cost of execution time and eventually are unable to parallel the accuracy of YA^14, 27^. Conforming to these findings, our results showed that OA achieved comparable *error rates*, whereas *performance time* was markedly increased. Although in line with the literature, this was a somewhat surprising finding, as we expected increased *error rates* for OA due to the substantial complexity of our paradigm. These results can be explained by the performance of a top-performing subgroup of OA. Within this group (N=10), OA paralleled and occasionally outperformed the accuracy of YA drawing the overall group mean of *error rates* towards the mean of YA. The main effect of hand and the interaction between complexity and hand for the factor *performance time* suggest that, regardless of age, subjects had difficulty to tap a new number sequence with the non-dominant left hand.

### 3.2 Increased and spatially extended cortical activity in older adults

Statistical analysis of ERP data consistently demonstrated enhanced positive potentials in older adults compared to young adults. In both experimental conditions, OA showed these enhanced positive potentials in distributed clusters located mainly above the hemisphere contralateral to the hand tapping a new sequence. This finding is in concordance with a study by Wu and Hallett who found that OA showed an increased BOLD-signal, interpreted as enhanced cortical activity, and activated additional sources while performing complex automatic movements at the same accuracy as YA^2^. Since the present analysis included only correct trials, our findings support the hypothesis that older adults rely on the recruitment of spatially extended brain areas to match the performance of YA during complex movements.

Analysis of the stimulus-locked ERP components showed no differences between both groups during early phases of the stimulus analysis, whereas pronounced differences could be revealed for the later phase. This is in accordance with previous ERP studies showing that age-related differences particularly emerge during later cognitive components, i.e. the P300, over central and parietal regions^14, 26^. In the present study, YA showed a marked positive deflection at 450 ms post “go”-stimulus, which can be interpreted as P300 as the deflection lies within the typical P300 latency range (250-500 ms)^28^. In contrast, a sustained positive deflection was observed in OA. The P300 is an ERP component elicited when attention has to be drawn to integrate sensory information with working memory^29^. Thus, the P300 seems to reflect the processing of a stimulus as well as the overall level of attention^28^. The amplitude of the P300 has been associated with memory processes but it is also thought to represent the amount of attentional resources engaged^28^. Although evidence exists that P300 amplitude declines with age when elicited by visual or auditory stimuli^30^, Sterr and Dean showed increased P300 amplitudes in OA during unimanual movement preparation^16^. Our results support the notion, that amplitudes of later cognitive components of the ERP such as the P300 are increased in OA compared to YA during preparation of movements. We speculate that this increase in amplitude along with the sustained positivity represents the higher cognitive demand OA have to engage to match the performance of YA. This is also supported by our supplementary analysis in which we compared YA with low and high performing OA and could replicate our findings. In conclusion, our results extend the findings of Wu and Hallett as well as Sterr and Dean in that a broader activation profile as well as an increased amplitude of cognitive ERP components is also present during the preparation of bimanual movements.

### 3.3 Absence of M1 activity and hierarchy in premotor areas

In the present study, we found no evoked source activity in M1 in either of the two conditions or groups (cf. 5.8 Source Specification). Although subsets of neurons in the primary motor area are active during movement preparation^31^, there is evidence for gating mechanisms which inhibit M1 neurons during movement preparation to prevent premature action^32^. It is therefore likely, that we were not able to detect this complex interplay of inhibition and facilitation within M1 when averaging ERPs time-locked to the “go”-stimulus. By choosing to average time-locked to the “go”-stimulus, we focused on the cognitive and preparatory part of the movement and not execution per se. Furthermore, the electrophysiological activity time-locked to the movement itself is very likely to vary inter-individually in relation to the “go”-stimulus and therefore averages out over several trials.

Random effects Bayesian model comparison revealed evidence for a top-down influence from lPM to SMA during preparation of complex bimanual movements. Numerous studies have shown the importance of both supplementary motor area^20, 33^ and premotor cortex (PM)^33, 34^ for the coordination of bimanual movements. Clear evidence for a hierarchy between SMA and PM, however, could not be detected in anatomical studies^35^. On the other hand, studies investigating the task-related involvement of both areas showed that SMA and PM are of importance for different subsets of bimanual movements. Here, SMA seems to be responsible for the temporal sequencing of movements and the parallel coding of actions of multiple effectors^20^. By contrast, the premotor cortex is engaged when movements depend on external stimuli and becomes particularly active when task complexity rises^34^. Since our paradigm involved complex bimanual movements an increased involvement of PM seems to be likely. In conjunction with the studies named above, our results support the notion that the bimanual coordination network is not a rigid entity. Rather, it is a dynamic system in which top-down control can change when adapting to task demands^20^.

### 3.4 Inhibitory feedback from premotor to prefrontal areas might represent conveyance of efference copies

Bayesian model averages of the A-matrix parameters common across all subjects consistently showed inhibitory bottom-up connectivity from premotor to prefrontal areas. It is widely acknowledged that the prediction of a movement-outcome relies on so-called forward models, which are simulations of the motor system about the consequences of an action^36, 37^. The forward model uses a copy of the motor command, the efference copy, which is thought to be generated in the supplementary motor area as well as the premotor cortex^38, 39^. We speculate that the bottom-up feedback from premotor areas could represent the conveyance of efference copies providing information about the expected consequences of movements. Since the PFC is engaged in online movement monitoring and online error correction^40, 41^, the inhibition of PFC could represent a reduced need for top-down error correction when the motor system expects a correct execution (only correct trials were used for analysis). This notion is supported by the findings that reduced inhibition of PFC from lPM was associated with higher error rates, whereby reduced inhibition of PFC from SMA was associated with higher performance time. This interpretation requires further investigation.

### 3.5 The distinct role of the PFC in increased intra- and interhemispheric coupling in older adults

There is a dominant view that the prefrontal cortex plays an eminent role in the cognitive control of action^42^. It has been suggested that activity in the dorsolateral prefrontal cortex reflects “attention to action” and that specific subregions within the PFC play a crucial role in online movement monitoring and working memory^40, 41, 43^. Furthermore, functions, ascribed to the PFC, seem to show a hemispheric predominance. Left PFC is supposed to be implicated in task setting and switching, whereas right PFC seems to be particularly involved in online movement monitoring and memory retrieval^41^. In the present study, high levels of attention were required to perform the complex bimanual movements correctly. In fact, two previous studies from our group as well as another study from Meister et al. demonstrated that precise prefrontal to premotor coupling is important for the execution of complex bimanual movements and that distinct coupling changes are associated with healthy ageing and Parkinson’s disease^4, 44, 45^. The influence of prefrontal to premotor coupling during movement planning, however, has not been investigated hitherto and we will discuss these interactions in the subsequent sections.

#### 3.5.1 Increased interhemispheric coupling in older adults

Converging evidence from TMS studies suggests that interhemispheric inhibition (IHI) between both primary motor cortices as well as premotor and primary motor areas is reduced (i.e. more facilitatory) in OA compared to YA^17, 46, 47^. Interhemispheric inhibition during movement preparation, however, is essential for bimanual movements^19^. In this study, Bayesian Model Averaging (BMA) revealed that interhemispheric connections from PFC to contralateral lPM were excitatory in OA whereas they were inhibitory in YA. These results conform to a study that has shown PFC over-activation during movement preparation of unimanual movements^14^. The increased activity in prefrontal areas, interpreted as enhanced reliance on cognitive control in OA, was seen as a compensatory mechanism as it was associated with better motor performance. Our results, however, allow for a more detailed interpretation of prefrontal influence during movement preparation as excitatory interhemispheric PFC to lPM connectivity was associated with poorer motor performance, arguing for de-differentiation in the prefrontal-premotor system. On the other hand, right-sided intrahemispheric connectivity was associated with better motor performance, giving evidence for a compensatory mechanism within this system. Interestingly, older adults in the top-performing group, who matched the performance of young adults, showed excitatory left PFC to right lPM coupling that was lower than the coupling of older adults in the low-performing group. Therefore, we argue that weak excitatory top-down influence from prefrontal to premotor areas can serve as a compensatory mechanism but impairs bimanual movements as excitatory top-down control increases. The importance of interhemispheric prefrontal to premotor top-down control during movement preparation is underscored by the finding that leave-one-out cross validation predicted the group association based on the two interhemispheric connections. In summary, we show, for the first time, that excitatory top-down control from prefrontal to lateral premotor areas is present in older adults during movement preparation. As interhemispheric inhibition during preparation of bimanual movements is important, our results suggest that age-related increases in prefrontal top-down control on contralateral premotor areas is impedimental for bimanual movements.

#### 3.5.2 Increased intrahemispheric coupling in older adults

The age-related prefrontal over-activation described in imaging studies was also detected during unimanual movement preparation in an electrophysiological analysis. Employing a reaction time task Berchicci and colleagues could demonstrate increased prefrontal activation in OA during a simple as well as a complex condition which was related to increased movement accuracy^14^. The increased right-sided prefrontal to premotor coupling in OA in the present study can be interpreted as elevated prefrontal activation but further delineates how prefrontal activity influences motor output. Importantly, the described coupling was associated with reduced *error rates* but increased *performance time*. Evidence exists that rising complexity of bimanual movements intensifies activity of premotor cortex and that this increased activity, particularly of the right premotor cortex, is important for bimanual control^34, 48^. Our results show that the increased recruitment of the lateral premotor cortex in OA is mediated via top-down control of the right PFC. This increased recruitment was associated with reduced errors whereas increased activation of SMA was correlated with increased performance time. We suppose that the excitatory top-down influences from right PFC to SMA and lPM are an important underlying factor for the frequently described phenomenon that OA can match the performance of YA at the cost of reduced movement speed.

### 3.6 Differences between the performance of both hands

Using Bayesian model averages of B-matrix parameters we could investigate the neural dynamics underlying the performance difference between both hands, which was revealed in our behavioural analysis. Here, in common across subjects, left PFC to SMA coupling was weaker in condition 1 compared to condition 2 which was associated with increased *performance time*. This association suggests that performance differences between hands, i.e. longer *performance time* for trials in which the new sequences had to be tapped with the non-dominant left hand, is at least partly due to a reduced left PFC to SMA coupling. It is noteworthy, that left prefrontal cortex is particularly important for task switching. In this regard, several studies could link left PFC activity to tasks in which subjects had to constantly update and switch the task set^49, 50^. In the present study, subjects had to tap a new sequence in every trial, requiring participants to update the task continuously. This seemed to pose a difficulty for young and older adults when the new sequence was tapped with the non-dominant left hand. The associated reduction in left PFC to SMA connectivity may represent reduced left prefrontal control and seems to be an important underlying factor for performance differences between hands.

### 3.7 Limitations

The present study has several limitations. First, we pursued a region of interest approach and included predefined cortical sources based on their activity in previous studies. Although it is likely that additional cortical and subcortical sources such as the basal ganglia, the cerebellum, and the anterior cingulate cortex play a role in bimanual movement preparation, we did not include these areas to reduce data dimensionality and to account for the limited capability of electroencephalography to model subcortical sources. Second, Bayesian model selection chooses the model that describes the data best within a given model space^51^. This entails that models with an architecture not incorporated in our model space might be equally or more probable. Third, the cross-sectional design of our study enabled us to investigate age-related differences in effective connectivity. To investigate age-related and subject specific changes in effective connectivity, longitudinal studies are necessary.

## 4. Conclusion

In the present study, we investigated the influence of healthy ageing on the neural dynamics within the prefrontal-premotor network during the preparation of a complex bimanual task. We could show that distinct connectivity profiles were common across age-groups, whereas others differed between young and older adults. Hence, several conclusions can be drawn from our results. First, OA seem to rely on an increased recruitment of spatially extended areas and engage more attentional resources as represented by more positive amplitudes of cognitive ERP-components to match the performance of YA. Second, correct and rapid execution of bimanual movements appears to require inhibitory bottom-up connections from prefrontal to premotor areas, which might represent the conveyance of efference copies. Third, interhemispheric inhibition from prefrontal to premotor areas was reduced in OA, whereby weak excitatory interhemispheric connectivity might represent a compensatory mechanism in top-performing OA but eventually leads to poorer performance as excitatory influences increase. Fourth, increased intrahemispheric right PFC to premotor coupling in OA was associated with increased performance time but reduced error rates, suggesting an important involvement in the frequently seen phenomenon that OA are able to match the performance of YA but require more time to perform the movement. Fifth, participants had difficulties when performing a new sequence with the non-dominant left hand, which was associated with reduced left PFC to SMA coupling. Hence, reduced left PFC influence might underlie performance differences between right and left hand when constant task monitoring and updating is required.

## 5. Materials and Methods

### 5.1 Participants

28 young and 32 old healthy volunteers participated in this study upon written informed consent. Following EEG-preprocessing, the datasets of 19 young (mean age ± SD: 25.79 ± 4.29 years; 5 female) and 20 old (mean age ± SD: 59.65 ± 6.82 years; 8 female) subjects were included for further analysis (see “Data Acquisition” and “EEG Preprocessing”). All participants were right-handed according to the Edinburgh Handedness Inventory, did not play an instrument for more than five hours per month, and had normal neuropsychological test scores (Mini-Mental State Examination, cut-off ≤ 25; DemTect, cut-off ≤ 13, and Beck’s Depression Inventory, cut-off ≥ 13). The experimental procedures were conducted in accordance with the Declaration of Helsinki and approved by the local ethics committee (study number: 13-394). The same data set has been previously published in another study^4^.

### 5.2 Experimental Paradigm

Experimental procedures were performed in an electrically and acoustically shielded room. Participants were seated comfortably in front of a computer screen with their fingers placed on a response pad (Cedrus, San Pedro, USA). The response pad consisted of eight buttons, four buttons for each hand, whereby a number as well as a corresponding button were allocated to each finger (1 for left and right thumb; 2 for left and right index finger; 3 for left and right middle finger and 4 for left and right ring finger). A comprehensive description of the experimental conditions and the behavioural paradigm is found in Loehrer and Nettersheim et al.^4^. In short, participants were instructed to learn a sequence of four button presses for one hand (e.g. left hand 1, 2, 3, 4). The learned sequence had to be executed with the respective hand while, simultaneously, the other hand had to tap a different sequence that was presented on the screen (see Figure 1.A). Participants were instructed to focus on correct trial execution rather than speed. Learned sequences were performed with the right hand (condition 1) and left hand (condition 2), whereas the starting hand was counter balanced across subjects. The behavioral task comprised six different tapping sequences of varying complexity which had to be learned and each of which had to be executed 20 times for both hands. At the beginning of each trial, the upcoming sequence of numbers was shown in red and signaled the subject to prepare for movement initiation. The switch from red numbers to green numbers indicated to start tapping (“go”-signal). A red cross marked the end of one trial, followed by a short break of 5 seconds. Trials in which subjects pressed the learned as well as the new sequences in the correct order were included for further analysis only. In addition to the condition analyzed here, participants completed a task in which they had to tap the same sequences with both hands.

**Figure 6.**
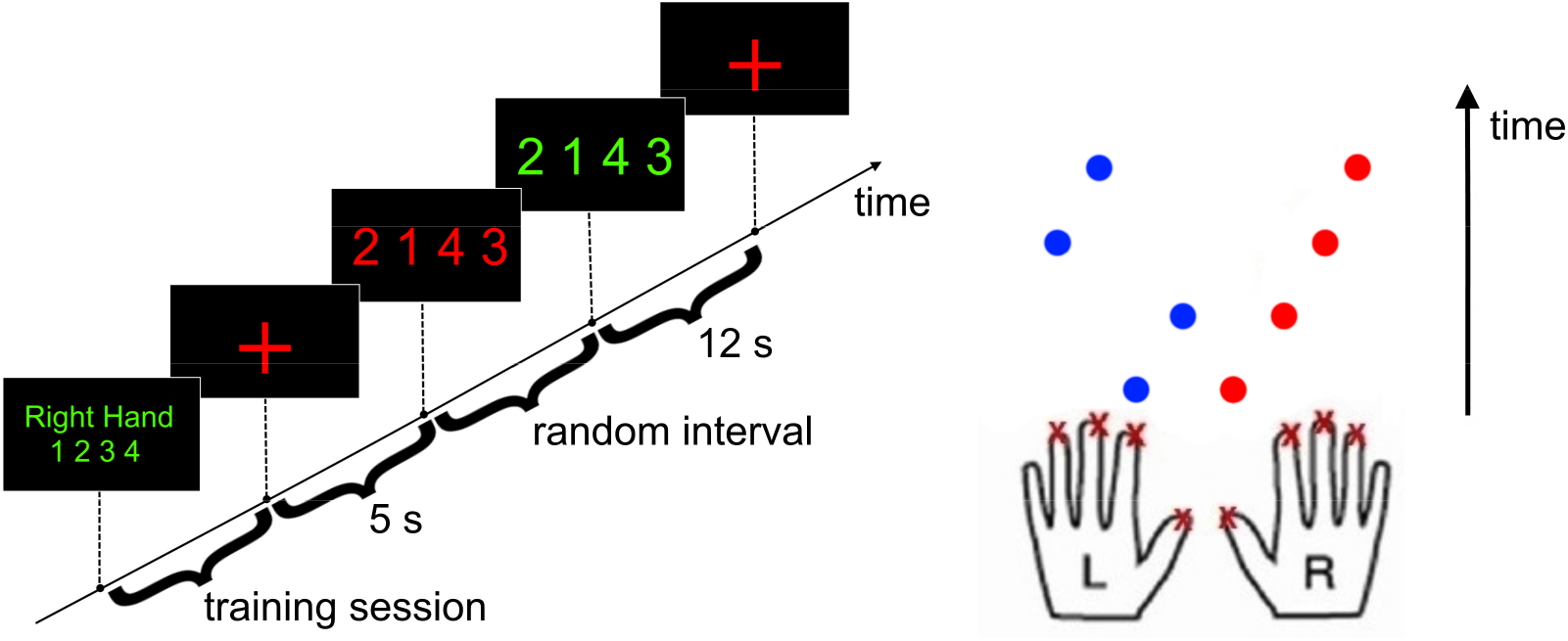
Instructions presented on a screen (left) and demanded button presses (right) in chronological order. In this example, participants had to learn a sequence for their right hand (condition 1) and tap a new sequence with their left hand. Hence, the first demanded pair of buttons was right thumb and left index finger.

### 5.3 Analysis of Behavioural Data

*Error rate* denotes the ratio of correct trials to overall trials. *Performance time* denotes the time between the go-signal and the last button press. Statistical analysis was performed using SPSS 22.0 (IBM, Armonk, USA) for Windows 10. The Shapiro-Wilk test revealed a non-Gaussian distribution for values of *error rate* and *performance time*, which prompted us to employ square root transformed data for further analysis. We used independent samples t-tests (two-tailed) to investigate overall differences between young (n=19) and older adults (n=20) in *error rate* and *performance time*. To evaluate differences in top performing old subjects compared to low performers and young adults, we conducted a median split for the variable *error rate. Error rate* and *performance time* were entered separately into a mixed-design ANOVA with the repeated measures “complexity” (levels 1-6) and “hand” (learned sequence performed with left or right hand) as well as the independent variable “group” (Young (n=19), Old – top performer (n=10), Old – low performer (n=10)) using Greenhouse-Geisser correction for non-sphericity when appropriate. Post-hoc Tukey tests were conducted comparing the three groups. We set all alpha-levels to 0.05. Effect sizes were calculated according to Cohen^52^.

### 5.4 Data Acquisition

EEG-data were acquired using a 128-channel system (Brain Products GmbH, Gilching, Germany) in an electrically shielded room. EEG-caps were placed upon subjects head and correct positioning was ensured by measuring the electrodes distances to nasion, inion, as well as left and right preauricular fiducials. After assuring that electrode impedances were below 10 kΩ, EEG-signals were recorded with Brainvision recorder (Brain Products GmbH, Gilching, Germany). Signals were amplified, band-pass filtered from 0.1 to 1000 Hz, and digitized at a sampling rate of 5000 Hz. Prior to EEG recordings, individual T1-weighted magnetic resonance images (MRI) of the head were acquired on a 3-Tesla Trio scanner (Siemens, Erlangen, Germany) using a 3D Modified Driven Equilibrium Fourier Transform sequence (repetition-time = 1930 ms, echo-time = 5.8 ms, flip-angle = 18°, slice-thickness = 1.25 mm). Three young and one older adult discontinued MRI acquisition and subsequent EEG recordings due to claustrophobia.

### 5.5 EEG Preprocessing

Data pre-processing was performed with BESA Research 6.0 (“brain electrical source analysis”, BESA, Gräfelfing, Germany). During offline analysis, the data was average-referenced and artefact-corrected. The amplitude threshold for excluding a channel from the analysis was below 0.07 μV, above 120 μV, or a gradient larger than 75 μV to adjacent channels. Noisy trials were excluded from further analysis. Eyeblink removal was performed with the automatic artefact removal tool provided by BESA (threshold for horizontal and vertical eye movements: 150 μV and 250 μV) or, if the method failed to remove all eyeblinks, with an additional independent component analysis (ICA). For calculating event-related activity, data was filtered (Butterworth filter: 0.53 – 200 Hz, notch filter: 50 Hz), downsampled to 1000 Hz, and averaged time-locked to the presentation of the green numbers (“go”-stimulus) for correctly performed trials. Subjects with too many noisy channels (> 60), too little correct trials (< 10), or without clear evoked activity were excluded from the analysis (6 young subjects, 11 old subjects). For DCM analysis, event related potential data was transformed to SPM data format (SPM12, update revision number 7487, Wellcome Trust Centre for Neuroimaging, London, UK), bandpass filtered between 1 and 48 Hz and downsampled to 250 Hz.

### 5.6 Statistical Analysis of Event-Related Potentials

Statistical analysis of event related potentials (ERP) was based on permutation testing to avoid alpha-error accumulation and was performed with BESA statistics 1.0. As a first step, for every sample of the ERPs a two-sided unpaired t-Test was calculated and significant differences between groups were defined (threshold: α = .05). Clusters were specified by selecting connected data points on the basis of temporal and spatial adjacency^53^. Subsequently, for each cluster, a cluster value was derived consisting of the sum of all t-values in the cluster. Then, the data of subjects were permutated, i.e. systematically interchanged. For each of the permutations a new t-test was computed and a new cluster-value derived. Thereby, a new distribution of cluster-values was obtained for each of the initial clusters. Based on the new cluster values resulting from the permuted data, it was possible to determine the significance of the initial cluster-value. If less than 5 % of the new cluster values were larger than the initial cluster, the initial cluster was considered significant with a p-value of .05. Effect sizes were calculated according to Cohen^52^.

### 5.7 Dynamic Causal Modelling

Dynamic causal modelling is a statistical approach to model the underlying neuronal architecture that gives rise to measured hemodynamic or electromagnetic time series data^22^. Employing neural mass models and Bayesian statistics, DCM allows one to infer the effective connectivity between brain regions and how these parameters are influenced by experimental factors^22^. Here, we used DCM for evoked responses to model the neuronal dynamics underlying movement preparation of a bimanual task in young and old subjects.

DCM for evoked responses rests on neural mass models with three subpopulations of neurons, as described by Jansen and Rit, to model source activity^21, 54^. Within this model, each subpopulation is assigned to one of three cortical layers: inhibitory interneurons in the supra-granular layer, excitatory neurons (spiny stellate cells) in the granular layer, and deep pyramidal cells in the infra-granular layer. Within a source, the three different layers are connected via excitatory and inhibitory interneurons. Connectivity between sources is established via pyramidal cells in the infra-granular layer. Here, bottom-up processing (forward connections) is represented by connections terminating in the granular layer, whereas top-down processing (backward connections) is mediated via connections terminating in the infra- and supra-granular layers. Lateral connections originate in the infra-granular layer also and terminate in all layers^21^. Experimental perturbations of the connectivity within a network is implemented via an exogenous input *u*, which has the same characteristics as a forward connection, hence terminating in the granular layer^21^. This architecture is based on the hierarchical organization described in Fellemann and Van Essen and allows one to build models with multiple interconnected sources^21, 35^.

The response of each subpopulation to an input is characterized by a set of differential equations. These equations describe the average response of a subpopulation in terms of firing rate-dependent current and voltage changes^21, 55^. The change in neuronal state of a source (*ẋ*) represents the average depolarization of pyramidal cells and can be described as a function of input (*u*) and its previous state (*x*):

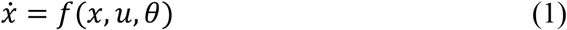

where *θ* represents the parameters and modulations of forward, backward, and lateral connections. These parameters are also called “hidden neuronal states” and have to be estimated. An electromagnetic forward model is used to estimate the change of neuronal state of a source (*x*_0_) based on a measured ERP signal (*y*):

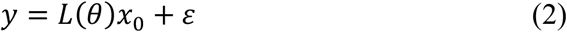

where *L*(*θ*) represents the parameterized lead field and *ε* observation noise (equations (1) and (2) are adopted from Garrido et al.^55, 56^). The combination of neural mass models with a spatiotemporal forward model results in a generative model that describes how model parameters and perturbations give rise to the measured ERP signal^22^.

Bayesian inversion of this generative model minimizes the Variational Free Energy under the Laplace Approximation^56, 57^. The employed procedure can be regarded as an Expectation-Maximization algorithm and provides an estimate of the posterior mean values of the model parameters *θ* and the log-model evidence. This model evidence is composed of the accuracy of a model, describing the data fit, and a complexity term, which penalizes models with increasing numbers of parameters^56, 58^. The model evidence can then be used to compare different models^51^.

### 5.8 Source Specification

In the framework of DCM, the specification of sources can be made using prior knowledge of the relevant sources based on previous studies^59^. In this study, activity of cortical sources included in subsequent dynamic causal models was confirmed in a source reconstruction of ERPs using a multiple sparse priors algorithm^60^. First, we defined a core motor network based on the MNI coordinates reported by Haslinger et al. and Herz et al. which were used in previous studies^4, 44, 61, 62^. Sources included in this network were dorsolateral prefrontal Cortex (PFC; MNI coordinates: left: [-37 48 22], right: [37 48 22]), lateral premotor Cortex (lPM; MNI coordinates: left: [-52 14 42], right: [52 14 42]), supplementary motor area (SMA; MNI coordinates: [0 −5 70]), and primary motor cortex (M1; MNI coordinates: left: [-41 −26 66], right: [41 −26 66]). Due to the spatial adjacency of bilateral SMA and the poor spatial resolution of EEG, we included SMA as a single source.

To evaluate whether these sources were indeed active during bimanual task performance, individual MRIs were coregistered with EEG electrode locations and a BEM (boundary element model) head model, employing a cortical mesh with 8196 dipoles, was calculated. Source activity for correct trials was inverted for both groups at 0-758 ms post-stimulus using the multiple sparse priors algorithm as implemented in SPM^60^. Length of this time interval was defined by the significant difference in ERP magnitude between young and old subjects (see “Results”). T-statistics were used to contrast inversion results to a 100 ms prestimulus baseline using family wise error (FWE) correction thresholded at <.05 at the peak and cluster level. To account for the low spatial resolution of EEG, we allowed for a deviation of 10 mm from our ROIs^62^. We could not find consistent activity in M1, which prompted us to exclude this source from our model space (cf. 4.3 Absence of M1 activity and hierarchy in premotor areas). Activity was confirmed in our other ROIs during the respective time interval (see Supplementary Figure 4).

### 5.9 Model Specification

Based on evidence from anatomical studies, we assumed reciprocal connections between lPM and SMA^35^ as well as reciprocal connections between PFC and lPM and PFC and SMA^35^. This connectivity structure was applied to the interhemispheric connectivity between prefrontal and premotor areas also. With respect to the employed hierarchy, i.e. the nature of reciprocal connections, we refer to a study of Felleman and Van Essen^35^ that suggests ascending (i.e. forward) connections going from lPM to PFC, and descending (i.e. backward) connections from PFC to lPM. Regarding lPM and SMA, there is no evidence for hierarchical differences.

Therefore, three fully connected models with either forward, backward, or lateral connections between lPM and SMA were inverted per subject. Model evidence was compared using a random effects Bayesian model comparison (see “Results”)^63^. In addition, lateral connections were used for interconnecting left and right PFC as well as lPM of both hemispheres. The input to the network was set to the PFC of both hemispheres, assuming prefrontal areas to drive the motor output as described in Herz et al.^64^.

### 5.10 Implementation of DCM in this Study

Dynamic causal modelling for evoked potentials was used to infer effective connectivity between five sources in 39 subjects in this study. A forward model was calculated for each subject as stated above (see “Source Specification”). ERPs at 0-758 ms post-stimulus were fitted to a fully connected model with backward connections from lPM to SMA and forward connections from SMA to lPM. This interval was necessary to capture the significant differences between young and old subjects in ERPs in both conditions (see “Results”). Activity of cortical sources was modeled using the equivalent current dipole (ECD) method as implemented in SPM^56^. To infer the effect of age on the average connectivity over conditions and on the difference between hands, we contrasted both conditions (condition 1: learned sequence performed with the right hand; condition 2 learned sequence performed with the left hand). Reduction of data dimensionality was achieved by singular value decomposition and the first eight principal components were used for inversion, which corresponds to the SPM default settings (see Supplementary Figure 5 for the frequency modes of a representative patient). A bandwidth from 4 to 48 Hz was used to avoid a 50 Hz electric current artifact.

### 5.11 Parametric Empirical Bayes and Bayesian Model Reduction

Parametric Empirical Bayes (PEB) describes the inversion of hierarchical models in which the posterior density of model parameters are constrained by the posterior density of the hierarchy level above^23^. The priors, i.e. the constraints, used for this inversion are derived from empirical data – contrary to standard Bayesian methods, for which the prior distribution is fixed – and are therefore called empirical priors^23^. This procedure involves inversion of a parent model (e.g. a fully connected model) for every subject at the first level. The parameters of interest (e.g. connectivity parameters) are then taken to the second level (i.e. group level). Here, a Bayesian General Linear Model is specified by collating the mean and the covariance matrices of the parameters of interest as well as a design matrix containing experimental effects called regressors (e.g. group affiliation, age, handedness)^24^. By integrating the covariance matrix into the GLM, modelling the parameters of interest at the group level takes into account the individual differences of each parameter. This approach, therefore, differentiates between hypothesized group-level effects and uninteresting random effects^24^.

In an ensuing Bayesian Model Comparison, PEB models with combinations of different parameters of interest can be compared to identify the model best explaining the differences and commonalities due to the entered regressors. Here, large numbers of models can be compared using Bayesian Model Reduction. BMR derives the parameters and model evidence of nested PEB models by systematically removing one parameter or subsets of parameters, i.e. setting the prior mean of the respective parameter(s) to zero^23^. If one assumes that all nested models are equally probable a priori, BMR can be used to perform an automated search over the model space and prune away parameters that do not contribute to the model evidence (an approach that we pursued here)^24^.

In the present study, the coupling parameters were assessed within the PEB framework to characterize the commonalities between subjects and the differences due to age-group (YA: n=19, OA n=20), performance-group (top-performer (n=10)/low-performer (n=10), error rate, and performance time. In a first step, the A-matrix parameters, which represent the average effective connectivity across experimental conditions, were entered into a PEB model with the mean-centered regressors mentioned above. BMR was used to perform an exhaustive comparison of different PEB models as described above. Subsequently, mean connectivity parameter estimates were derived by averaging the parameters across models weighed by the models’ posterior probability, a process called Bayesian Model Averaging^65^. BMA was thresholded at 95 % posterior probability to focus the results on parameters with strong evidence of being non-zero^24^. This procedure was repeated for the B-matrix parameters, which - in our study - represent the effective connectivity necessary to model the difference between both conditions. It is noteworthy that the PEB model for the B-matrix parameters comprised age-group (YA: n=19, OA n=20), performance-group (top-performer (n=10)/low-performer (n=10), as well as the difference between both conditions in *error rate* and *performance time* as mean-centered regressors. PEB models with additional regressors (such as age or handedness) were not superior to the selected PEB model as revealed by free energy comparison. To disentangle the interaction between coupling strength, age-group, and performance parameters, coupling parameters were correlated with performance parameters partitioned by age-group using spearman-correlation. We controlled the false discovery rate using the Benjamini-Hochberg method to account for multiple comparisons and report corrected p-values^66^. In a last step, we assessed the predictive validity of estimated connectivity parameters using leave-one-out cross validation^24^. First, we defined connections that were most relevant for age-group association by employing the mutual information quotient (MIQ) as described in Weber^67^. Subsequently, we sought to ascertain whether the effect size of the selected connectivity parameters was large enough to predict a subject’s group affiliation.

## 7. Acknowledgement

The authors would like to thank the participants for their active engagement in this study.

## 8. Author contributions

Study design: PAL, FSN, LT, and IW; Conceptualization: PAL and IW; Methodology: PAL FSN, FJ, CO, and IW; Formal analysis: PAL, FSN, FJ and IW; Investigation: PAL, FSN, FJ, CO, MT, and IW; Resources: MT and LT; Writing - original draft: PAL; Writing - review & editing, PAL, FSN, CO, LT and IW; Supervision: LT

## 10. Data availability

The data that support the findings of this study are available upon reasonable request from the corresponding author (PAL). MRI data are not publicly available since they contain information that could compromise research participant privacy/consent. A source data file is provided for Figures 1.A, 1.B, 3, 4, 5, and Supplementary Figure 3.

## 11. Code availability

We used the commercial software BESA research 6.0 (BESA, Gräfelfing, Germany) for preprocessing of EEG data and BESA statistics 1.0 (BESA, Gräfelfing, Germany) for statistical analysis of ERP data. Analysis of behavioural data was performed using the commercial software SPSS 22.0 (IBM, Armonk, USA). Auxiliary scripts and code used for further analysis was written and developed in MATLAB (The MathWorks, Inc., MA, USA; commercial software), predominantly on version 2016b and 2018a, and are available at https://github.com/Loehrerph/BiSyT. For DCM analysis we used SPM12 (update revision number 7487, Wellcome Trust Centre for Neuroimaging, London, UK) freely available at https://www.fil.ion.ucl.ac.uk/spm/software/spm12/. The mutual information quotient feature selection algorithm is freely available within the NoLiTiA-toolbox at https://www.nolitia.com/download.html. Furthermore, the MATLAB plugin BrainNet Viewer, freely available at https://www.nitrc.org/projects/bnv/, was used to visualize the schematics in Figures 3 to 5.

---

**Supplementary Figure 1.**
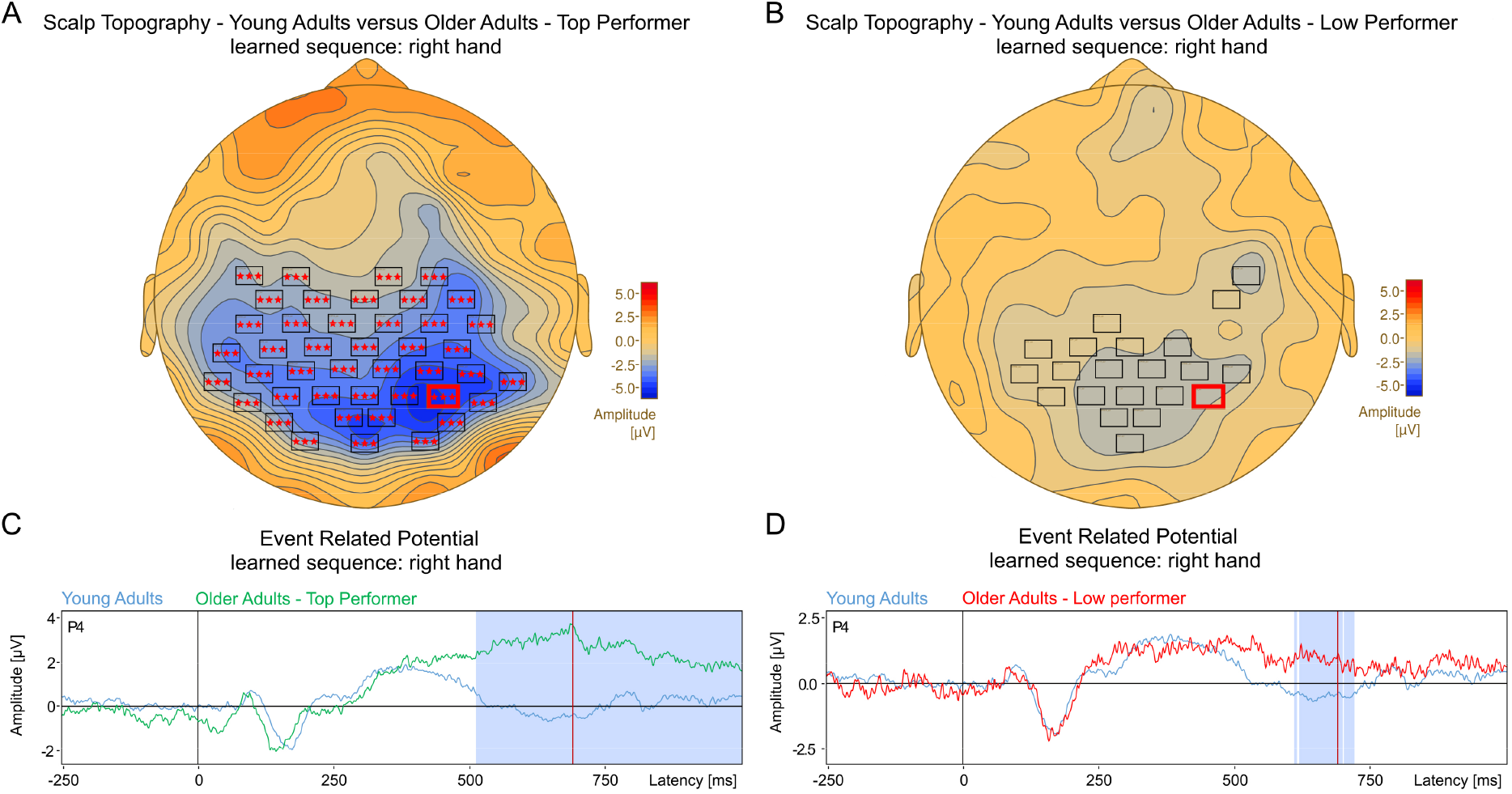
**S1.A.** Scalp topography comparing young adults and older adults – top performer. Topography is plotted for correctly performed trials at 688 ms after stimulus onset when the learned rule was performed with the right hand and the new number sequence with the left hand. The cluster with a p-value < .001 is denoted with three starlets (one-sided permutation test, Cohen’s d = 2.26, YA: n = 19, OA – top performer: n = 10). **B.** Scalp topography comparing young adults and older adults – low performer. Topography is plotted for correctly performed trials at 688 ms after stimulus onset when the learned rule was performed with the right hand and the new number sequence with the left hand. The cluster with a p-value of .057 is depicted (one-sided permutation test, Cohen’s d = 1.53, YA: n = 19, OA – low performer: n = 10). The red line in **A** and **B** shows the time point for which the scalp topography is plotted. **C.** ERP waveforms averaged time-locked to the “go”-stimulus: Grand-mean ERP of young adults (blue curve) and older adults – top performer (green curve) recorded with electrode P4 (right hand rule). Significant differences are shaded in light blue color. **D.** ERP waveforms averaged time locked to the “go”-stimulus: Grand-mean ERP of young adults (blue curve) and older adults – low performer (red curve) recorded with electrode P4 (right hand rule). Significant differences are shaded in light blue color.

**Supplementary Figure 2.**
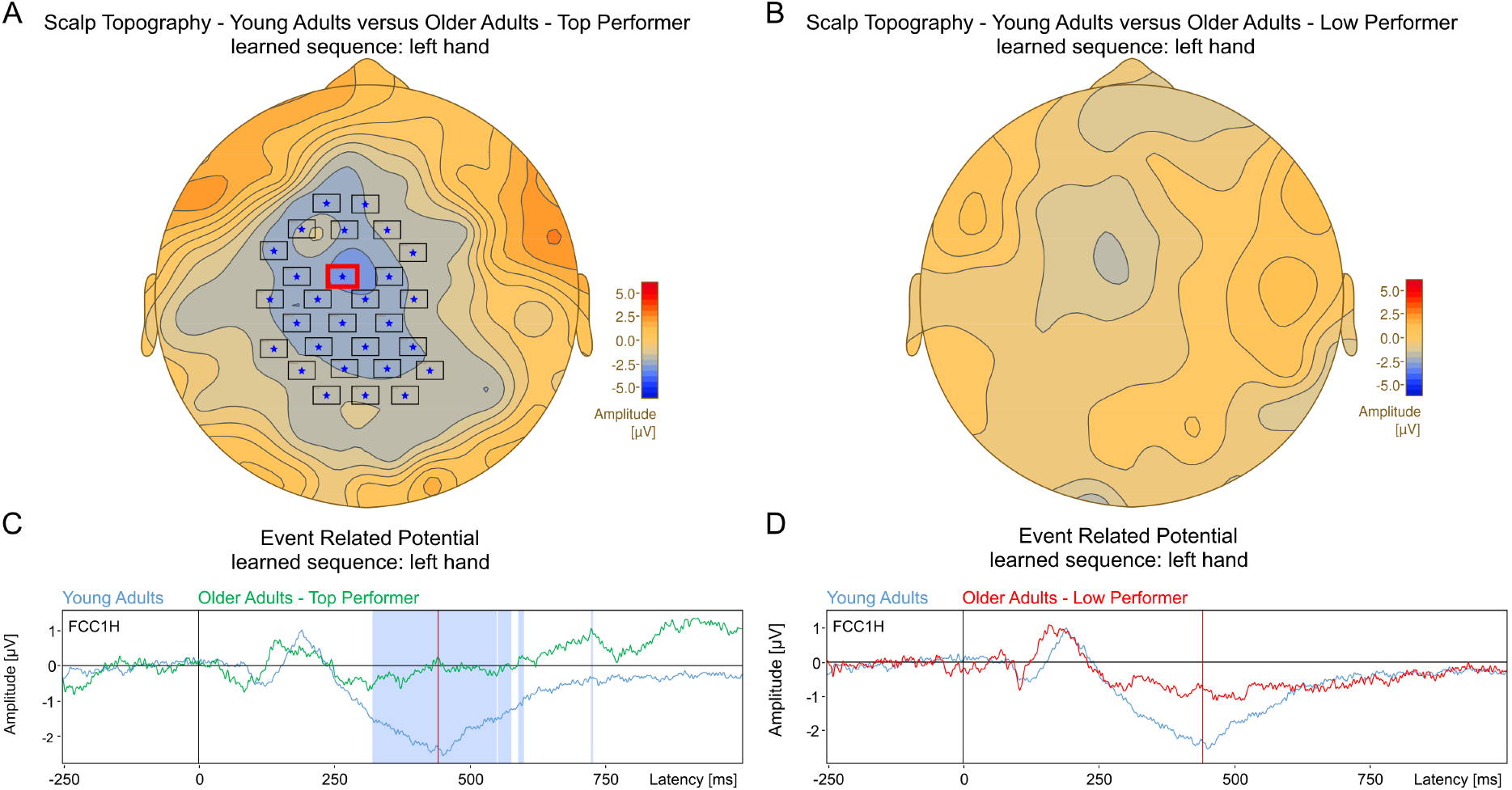
**S2.A.** Scalp topography comparing young adults and older adults – top performer. Topography is plotted for correctly performed trials at 440 ms after stimulus onset when the learned rule was performed with the left hand and the new number sequence with the right hand. The cluster with a p-value = .006 is denoted with a cross (one-sided permutation test, Cohen’s d = 1.8, YA: n = 19, OA – top performer: n = 10). **B.** Scalp topography comparing young adults and older adults – low performer. Topography is plotted for correctly performed trials at 440 ms after stimulus onset when the learned rule was performed with the left hand and the new number sequence with the right hand. No significant differences could be observed (cluster p-value = .786, one-sided permutation test, Cohen’s d = 1.18, YA: n = 19, OA – low performer: n = 10). The red line in **A** and **B** shows the time point for which the scalp topography is plotted. **C.** ERP waveforms averaged time-locked to the “go”-stimulus: Grand-mean ERP of young adults (blue curve) and older adults – top performer (green curve) recorded with electrode FCC1H (left hand rule). Significant differences are shaded in light blue color. **D.** ERP waveforms averaged time locked to the “go”-stimulus: Grand-mean ERP of young adults (blue curve) and older adults – low performer (red curve) recorded with electrode FCC1H (left hand rule).

**Supplementary Figure 3.**
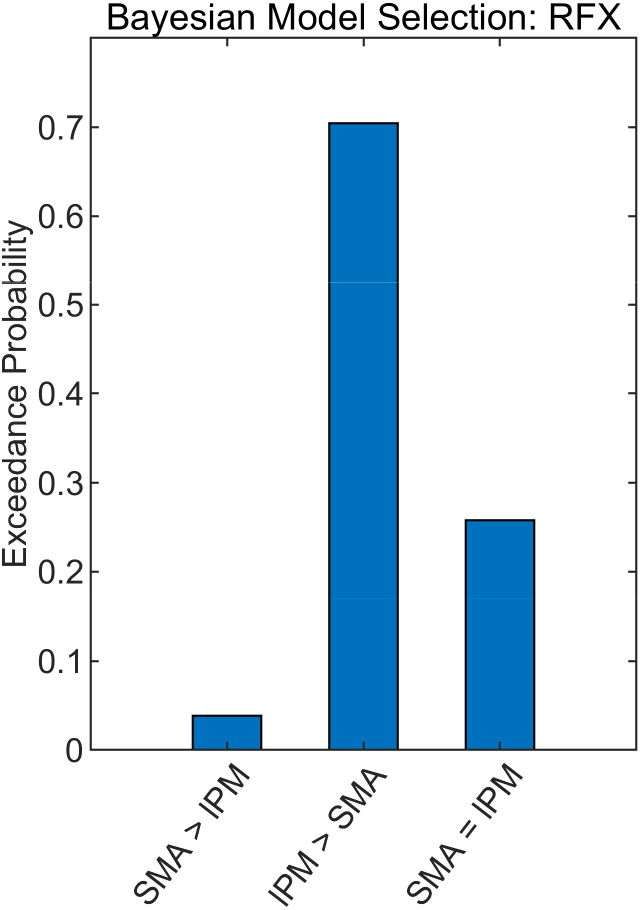
**S3.** Random effects Bayesian model comparison between three fully connected models, which differed in connectivity between SMA and lPM. SMA>lPM denotes the model with backward connections from SMA to lPM. lPM>SMA denotes the model with backward connections from lPM to SMA. SMA=lPM refers to the model with lateral connections between SMA and lPM. BMS revealed evidence in favor of the model comprising backward connections from lPM to SMA.

**Supplementary Figure 4.**
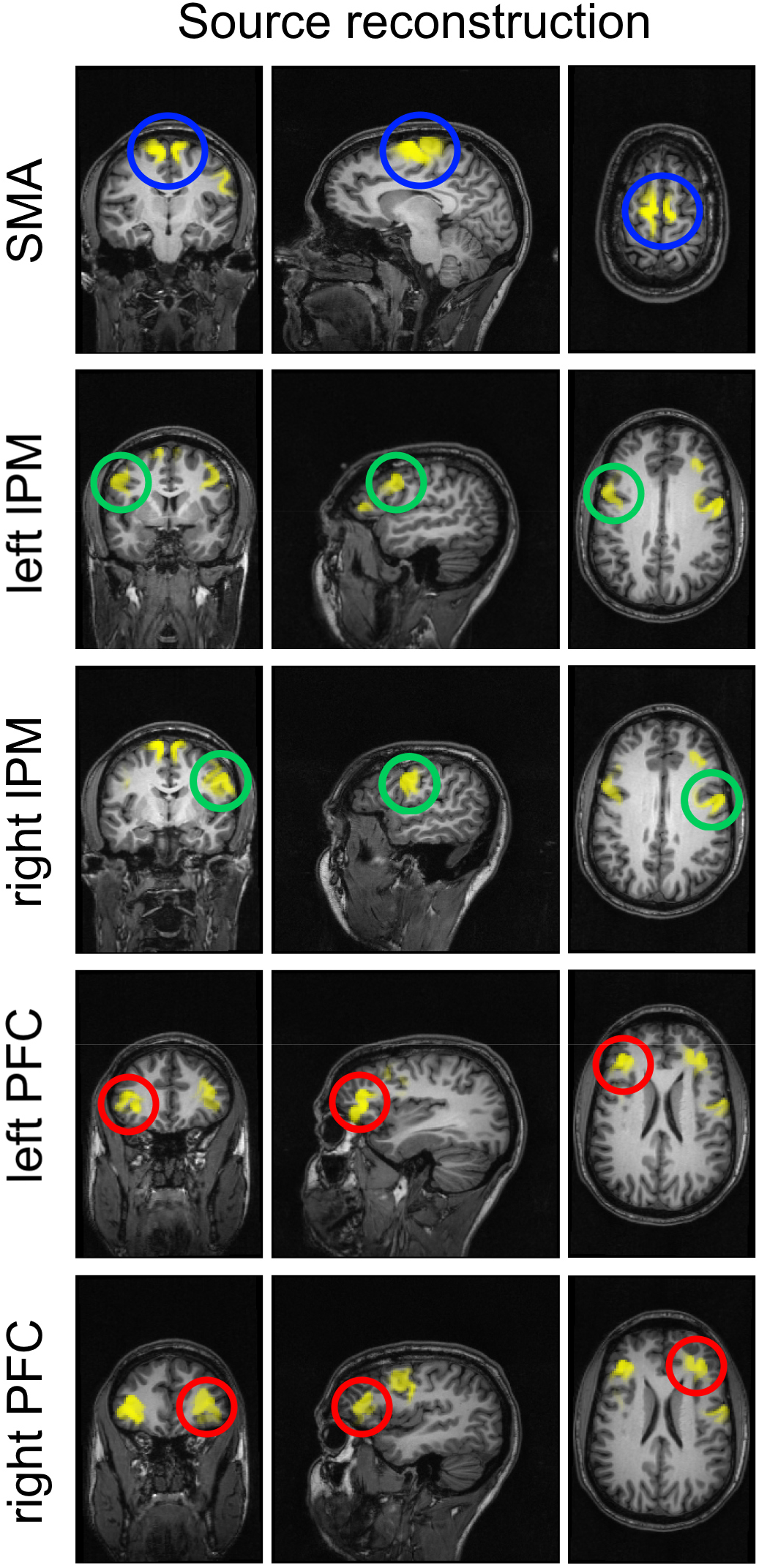
**S4.** Average source activity of condition 1 mapped onto a participants individual T1-weighted cranial MRI. Using the multiple sparse prior algorithm for correctly performed trials, we confirmed sources to be active for the time window of 0 to 758 ms.

**Supplementary Figure 5.**
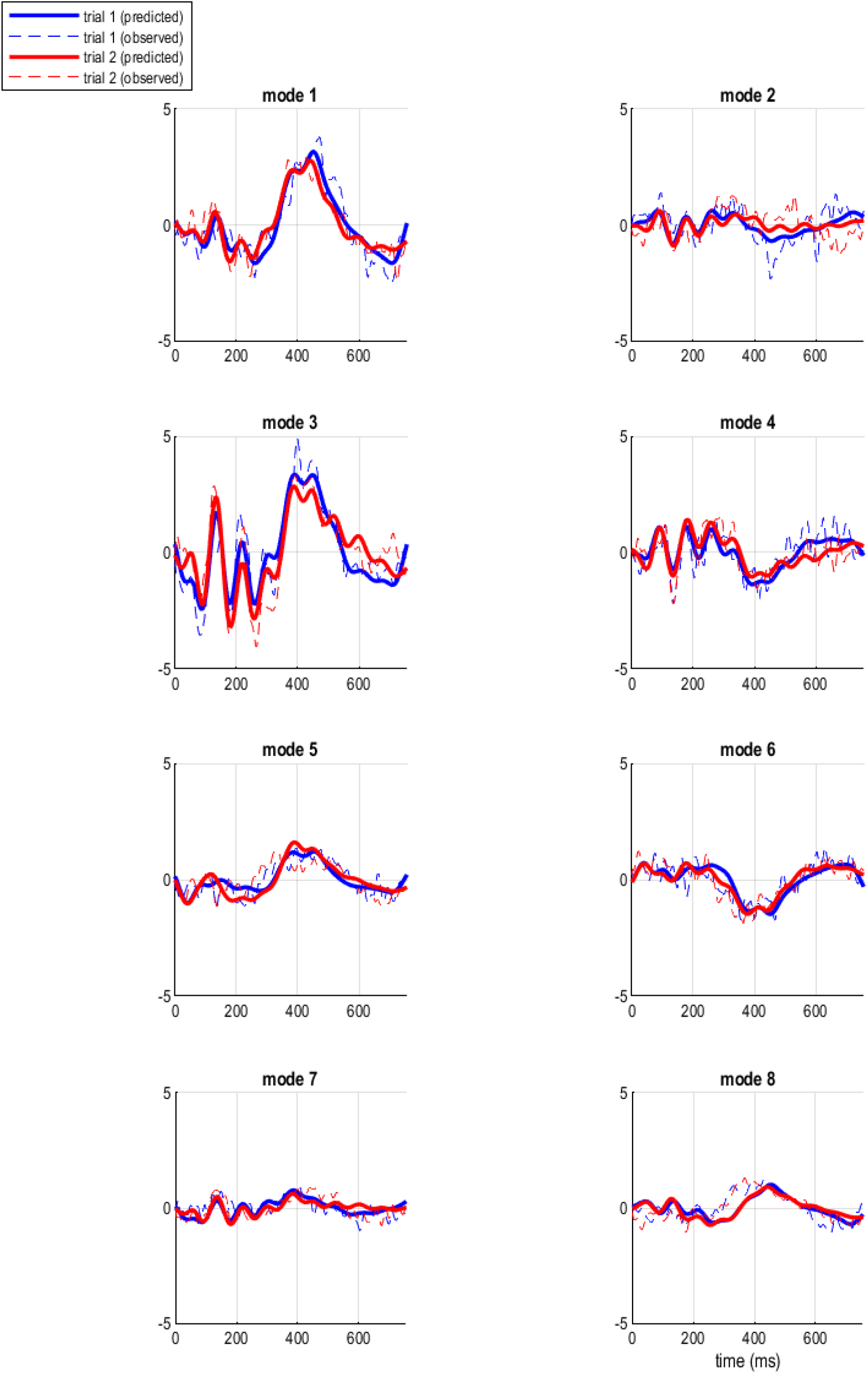
**S5.** Frequency modes of a representative patient. Power spectra were reduced in their dimensionality by singular value decomposition (SVD) to eight principal frequency modes.

